# Persistent Activation of Monocytes/Macrophages and Cell Senescence in SIV-Infected Macaques on ART

**DOI:** 10.1101/2025.11.05.686810

**Authors:** Yilin Chen, Xiaofeng Ding, Sonalika Ray, Siva Thirugnanam, Robert Blair, Ahmad Saied, Sergiy Sukhanov, Jay Kolls, Woong-Ki Kim, Patrice Delafontaine, Jay Rappaport, Xuebin Qin, Namita Rout

## Abstract

Despite effective viral suppression with antiretroviral therapy (ART), people living with HIV (PLWH) experience persistent inflammation, immune dysfunction, and premature onset of cardiovascular and aging-related comorbidities. To define the underlying mechanisms, we performed longitudinal transcriptomic profiling in peripheral blood mononuclear cells (PBMCs) from a cohort of simian immunodeficiency virus (SIV)-infected rhesus macaques spanning four key stages: pre-infection, acute infection, short-term ART, and long-term ART. Bulk RNA sequencing revealed dynamic immune remodeling across infection and treatment. Acute SIV infection induced robust antiviral and inflammatory programs, with upregulation of interferon-stimulated genes (ISGs), IL-27, JAK/STAT, and NF-κB signaling, coupled with suppression of T- and B-cell activation pathways. Short-term ART effectively reversed these transcriptional perturbations, restoring adaptive immune gene expression and reducing innate antiviral responses to near-baseline levels. In contrast, chronic SIV infection on long-term ART maintained viral suppression but was characterized by reactivation of innate immune pathways, including TLR2/TLR4/MYD88, NF-κB, and inflammasome (NLRP3/or NLRP12, caspase-1) signaling, along with sustained macrophage activation, platelet/coagulation signaling, and senescence-associated secretory phenotype. Protein analyses confirmed persistent CASPASE-1 and NF-κB activation in spleen tissue. Pathologic evaluation of a carotid artery from an SIV-infected, long-term ART– treated macaque revealed macrophage-rich plaques with p21⁺ senescent cells with intraluminal thrombus formation, recapitulating key features of HIV-associated atherogenesis. Together, these findings demonstrate that while ART normalizes acute infection–induced immune dysregulation, chronic SIV infection sustains a chronic, macrophage- and TLR-driven inflammatory state linked to vascular injury and aging process regardless of long-term suppression of viremia. Targeting inflammasome, NF-κB, and senescence pathways may mitigate non-AIDS comorbidities in PLWH.

## Introduction

People living with HIV (**PLWH**) are susceptible to inflammatory comorbidities despite effective antiretroviral therapy (**ART**) and experience premature onset of aging-related comorbidities such as cardiovascular disease (**CVD**). They are twice as likely to develop CVD (1) including atherosclerosis, a leading cause of morbidity and mortality in PLWH. PLWH also have an increased risk of thrombosis, which significantly contributes to atherosclerosis-associated CVD such as heart attack and stroke (2, 3). While HIV-induced chronic inflammation contributes to the accelerated aging process and CVD in PLWH (4), the causal mechanisms remain unclear (5). Chronic inflammation-associated biological aging , or inflammaging, is a significant contributor to CVD (6–8), and evidence suggests that chronic HIV infection accelerates inflammaging in PLWH (6, 9, 10). As the HIV-positive population ages, and with new diagnoses in older adults on the rise, understanding the complex interplay between HIV, aging, and inflammation has become increasingly important, particularly in the context of CVD pathogenesis. Elucidating the molecular and cellular mechanisms of HIV-induced acceleration of cardiovascular aging (6, 10) is crucial for the development of targeted interventions to prevent and treat HIV-associated CVD and other age-related diseases in PLWH (5, 11–14).

HIV infects CD4^+^ T lymphocytes and macrophages, triggering proinflammatory signaling cascades such as NLRP3 inflammasome-mediated caspase-1 (**NLRP3-CASP-1**) activation and the release of IL-1β and IL-18 (5, 15–19). Chronically activated monocytes/macrophages (MC/Mϕ) further aggravate HIV-associated atherosclerosis (**HIVAA**) (5, 11, 20–22). We (20, 21, 23–26) and others (27–31) have linked NLRP3-CASP-1 activation to HIVAA in PLWH and shown that macrophage-driven inflammasome activation promotes atherogenesis in mice; however, its role in accelerated aging and CVD requires further investigation in a physiologically relevant model (5, 27, 28, 32). Nonhuman primates (**NHPs**) naturally develop age-related CVD driven by risk factors similar to humans (e.g., dyslipidemia)(33, 34) and exhibit atherosclerotic lesions when fed atherogenic diet (**AD**)(34–39), providing a translational model to investigate HIV-associated CVD and accelerated aging. Prior studies using SIV-infected, CD8-lymphocyte-depleted macaques demonstrated CASPASE-1 and NF-κB activation in lymphoid tissues that persisted despite ART at 120 dpi (21). Whether such inflammatory signaling persists during chronic infection beyond this period remains unknown. Comprehensive longitudinal analysis of gene expression dynamics during HIV infection is not feasible in clinical settings, particularly because of the lack of access to pre-infection baseline samples and NHP studies to date have largely been limited to single-cell or bulk transcriptomic analyses of specific immune subsets at early infection with/without ART rather than systemic immune remodeling through chronic SIV infection under effective long-term ART (40, 41). Comprehensive profiling of global gene and pathway changes in the circulation specially in inflammaging pathways such as NLRP3–CASP1, NF-κB, and senescence pathways during an entire course of ART-suppressed HIV infection has yet to be defined in a physiologically relevant, non–CD8-depleted SIV infection model.

In this study, we established a longitudinal cohort of SIV-infected rhesus macaques to characterize transcriptomic remodeling in peripheral blood mononuclear cells (PBMCs) with infection and through long-term viral suppression for over a year with ART. PBMC’s bulk RNA sequencing (RNA-seq) was used to delineate persistent immune and inflammatory signatures linked to senescence and cardiovascular risk during chronic SIV infection under long-term ART. This approach revealed stage-specific immune remodeling, with acute infection driving broad antiviral and innate immune programs and ART exerting differential effects over time. While short-term therapy largely normalized adaptive and innate immune pathways, chronic SIV infection on long-term ART was associated with persistent immune activation and inflammatory signaling, including pathways linked to monocyte and macrophage activation, platelet activation, coagulation, and cellular senescence, all of which have been implicated in cardiovascular risk. Consistent with these systemic findings, we detected increased caspase-1 and NF-κB activation in spleen from SIV-infected rhesus macaques on long-term ART. Together, these findings delineate distinct immune transcriptomic trajectories during infection and therapy and suggest that durable viral suppression does not fully resolve pathogenic inflammatory programs that may contribute to HIV-associated comorbidities.

## Results

### Dynamic transcriptomic changes in SIV-infected RMs during acute infection and ART

We randomly selected 4 animals from a previous cohort (42) that had longitudinal PBMC samples collected at four key time-points: pre-SIV infection (day 0), 6 weeks post-SIV infection (acute SIV infection), after 2 months of short-term ART, and after one year of long-term ART, for RNA-seq analyses (Fig. 1a). This cohort effectively models the major stages of HIV infection in humans: 1) pre-infection, 2) acute HIV infection, 3) HIV infection on short-term ART treatment, and 4) chronic HIV infection on long-term ART treatment as seen in PLWH. Plasma viral loads peaked between days 7–14 post-infection and were rapidly suppressed following initiation of ART, remaining undetectable throughout the treatment phase (**Fig. 1b**). To comprehesively profile the dynamic transcriptomic changes in the PBMCs collected before and after SIV infection, with or without ART treatment, we conducted the bulk RNA-seq and performed six pairwise comparisons: 1) acute SIV-infection vs. baseline, 2) acute SIV-infection on short-term ART vs. acute SIV-infection, 3) acute SIV-infection on short-term ART vs baseline, 4) chronic SIV-infection on long-term ART vs acute SIV-infection, 5) chronic SIV-infection on long-term ART vs baseline, and 6) chronic SIV-infection on long-term ART vs chronic SIV-infection on short-term ART (**Fig. 1c**).

**Figure 1.**
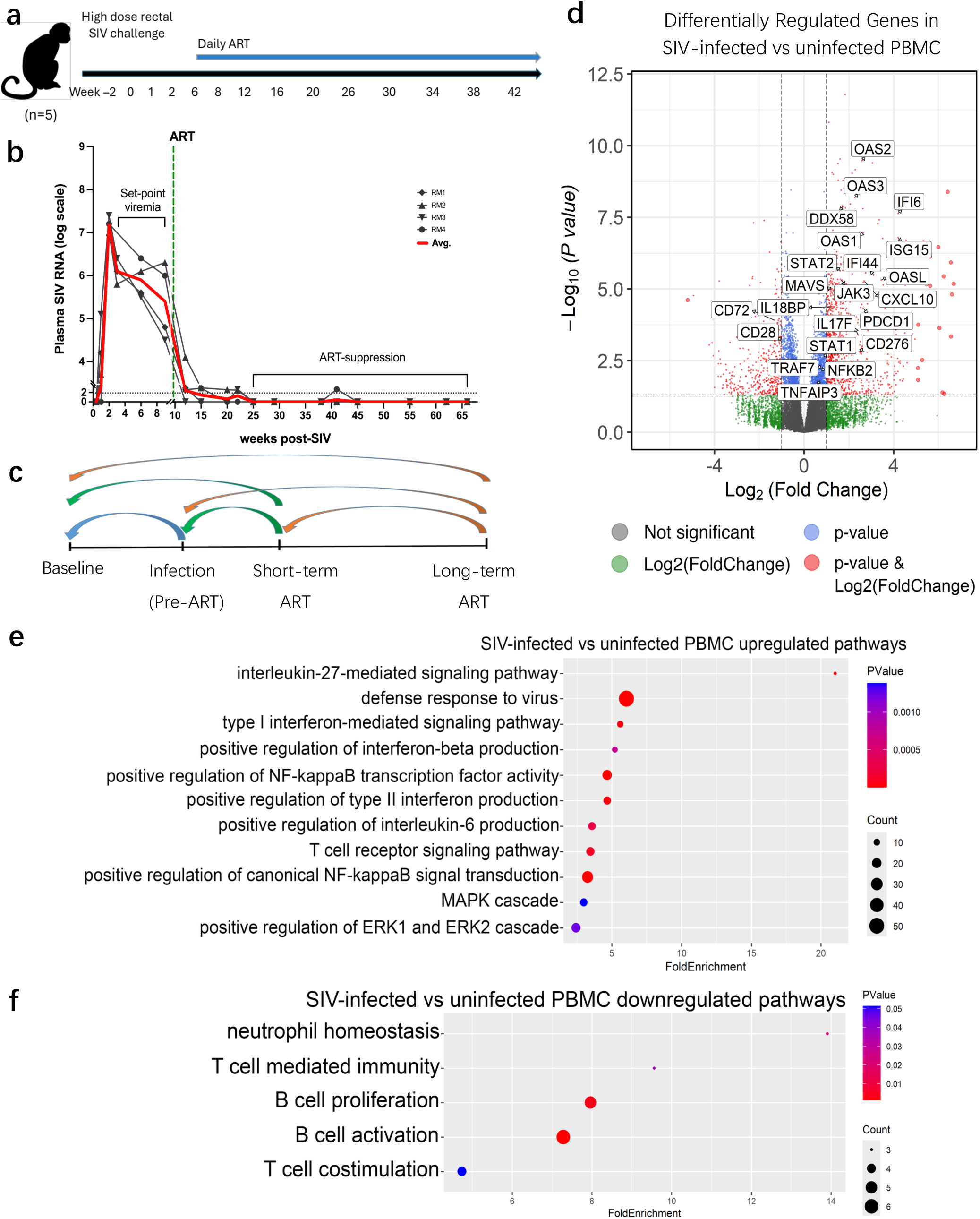
Acute SIV infection induces host innate antiviral programs while disrupting T- and B-cell activation pathways. **(a)** Timeline of SIV-infection and ART treatment in Rhesus Monkey cohort (n=5). **(b)** Plasma SIV RNA viral load for each animal. **(c)** Pairwise comparison design of bulk RNA-seq analysis. **(d)** Volcano plot showing differentially expressed genes (DEGs) in SIV-infected vs uninfected monkey. **(e-f)** Gene Ontology analysis of **(e)** upregulated and **(f)** downregulated DEGs in SIV-infected monkeys compared to uninfected PBMC.

### Acute SIV infection dramatically upregulates host antiviral innate immunity while dysregulating T cell and B cell activation pathways

Differential gene expression analysis of PBMCs revealed significant transcriptional changes between SIV-infected and uninfected animals (**Fig. 1d**). The volcano plot highlights several upregulated interferon-stimulated genes (ISGs)(e.g., *IFI44, ISG15, IFI6*), Type I interferon signaling responsive genes (*OAS1, OAS2, OAS3, OASL, STAT1, CXCL10*), antiviral signaling genes [e.g. RIG-I (*DDX58*), *MAVS*], and interleukin-signaling genes (e.g. *IL17F*, *IL18BP*) (**Fig. 1d**). Genes involved in JAK/STAT signaling (*JAK3, STAT2*), NF-kB signaling (e.g. *TNFAIP3, NFKB2*, and *TRAF7*) were also significantly upregulated, suggesting robust activation of immune pathways, mainly host innate immune response to viral infection (**Fig. 1d**). Of note, genes associated with T cell immune checkpoint *PDCD1* (PD-1), and *CD276* (B7-H3) were also upregulated in acute SIV-infection (**Fig. 1d**), suggesting activation of inhibitory signaling to counterbalance the robust ISG and antiviral responses. Consistently, genes associated with T cell co-stimulation (e.g.*CD28*), and B-cell activation (e.g. *CD72*) were downregulated (**Fig. 1d**). Togther, these results indicate that even though at the acute stage of SIV infection, innate antiviral programs were active, adaptive immunity is suppressed, potentially limiting effective cytotoxic T cell responses.

GO analysis further revealed functional pathway alterations. In line with volcano plot results, upregulated pathways in SIV-infected animals included those related to proinflammatory signaling pathway, including IL27-mediated signaling pathway, type I and type II interferon signaling, defense response to virus, IL6 production, IFNbeta signaling pathway, MAPK cascade, positive regulation of ERK1 and ERK2 cascade, and NF-κB signaling (**Fig. 1e**). In contrast, downregulated pathways includes T cell costimulation and B cell activation signaling pathways (**Fig. 1f**). These findings shows robust activation of antiviral and innate immune signaling pathways (43–45), and dysregulation of B-cell and T-cell activation(46, 47), in line with current understanding of HIV/SIV pathogenesis.

### Viral suppression with ART reduced host antiviral innate immune activation and restored adaptive immune gene expression to baseline levels

To further investigate the transcriptomics changes induced by ART treatment in SIV-infected rhesus macaques, we performed a pairwise comparison of the acute SIV infection on short-term ART vs the SIV-infected PBMC dataset. Differentially regulated genes in SIV-infected monkeys on short-term ART compared to SIV-infected only monkeys include upregulation of B cell activation signaling pathway genes (e.g, *CD79A, CD79B, CD40*, *CD72*) and T cell activation-related genes (e.g., *CD28*, *ICOS, DPP4*) compared to acute SIV-infection (**Fig. 2a**). Conversely, T cell inhibitory genes such as *PDCD1* (PD-1) and *CD276* (B7-H3) were downregulated, showing that short-term ART restored T cell and B cell activation in SIV-infected rhesus macaques (**Fig. 2a**). Additionally, other downregulated genes included IL27-related genes (e.g*.IL27*, *OAS1, OAS2, OAS3, OASL*), antiviral innate immune response [e.g. *DDX58* (RIG-I), *MAVS*)], ISGs (*ISG15, IFIT3, IFI6, IFI27)*, NF-kB signaling related genes (*IKBKG, IKBKE, TRAF7*), JAK/STAT signaling genes (*JAK3, STAT2*) (**Fig. 2a**). Notably, IL6, which is not only involved in activation of JAK/STAT, PI3k/AKT and MAPK signaling pathways, but also associated with senescnece signature, was also downregulated following short-term ART (**Fig. 2a**). Together, these results impliy that short-term ART treatment in SIV-infected macaques normalized the most significant gene expression changes induced by SIV infection (**Fig. 2a**).

**Figure 2.**
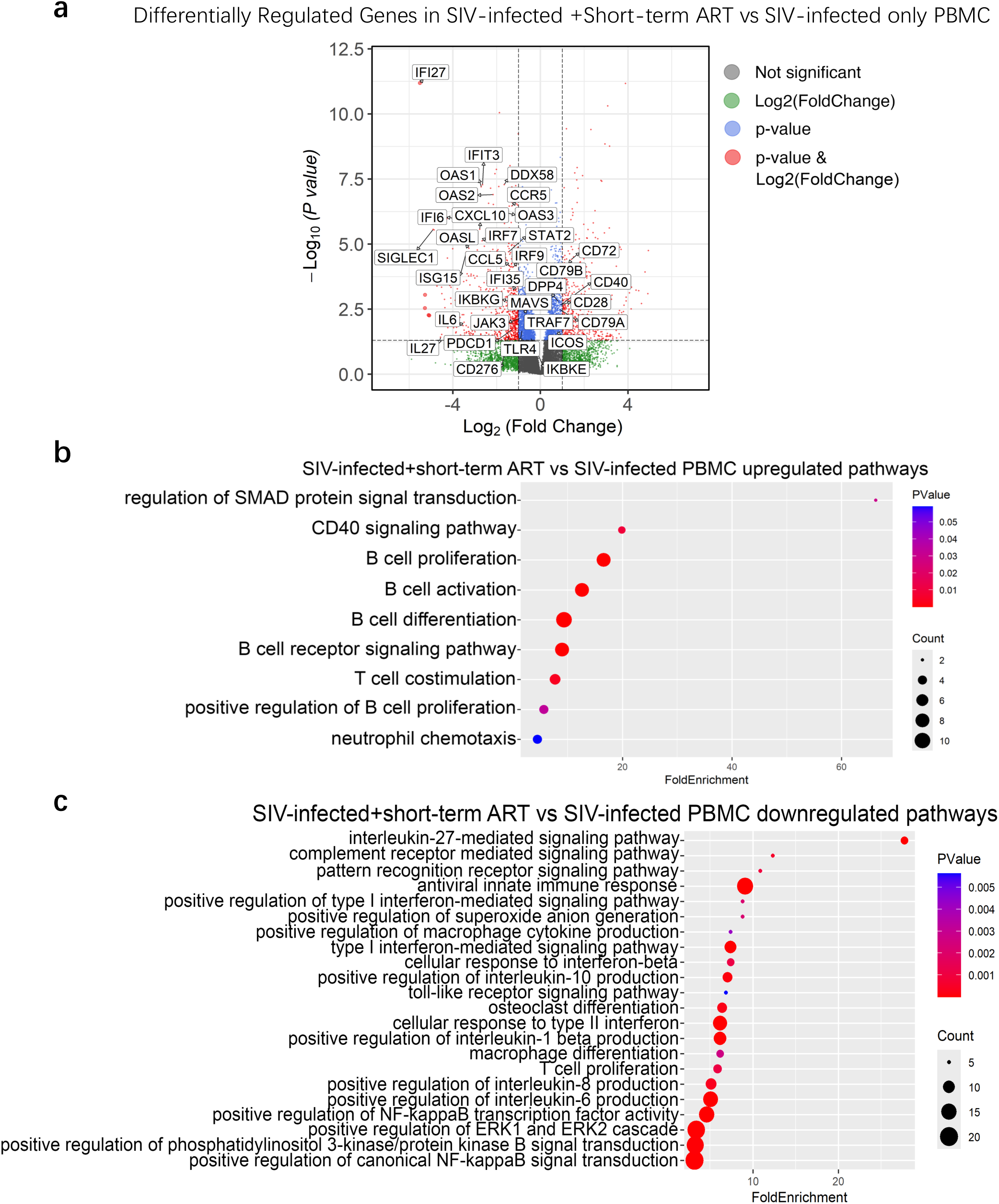
Short-term ART alleviates innate antiviral responses and restores adaptive immunity compared to acute SIV-infected macaques. **(a)** Volcano plot showing differentially expressed genes (DEGs) in SIV-infected + short-term ART vs SIV-infected only monkey PBMC. Gene Ontology analysis of **(b)** upregulated and **(c)** downregulated DEGs in PBMCs from SIV-infected + short-term ART macaques compared to that in SIV-infected macaques before ART initiation.

GO functional analysis on the differentially expressed genes (DEGs) in SIV-infection on short-term ART vs SIV-infected only dataset further confirms the trends seen in the volcano plot. Upregulated GO terms in SIV-infection on short-term ART include B cell activation, T cell costimulation, and CD40 signaling pathways, showing that short-term ART restores T cell and B cell activation towards immune homeostasis (**Fig. 2b**). Downregulated GO terms include IL27-signaling pathway, antiviral innate immune response, IL6 production, NF-kB signaling, and type I/II IFN signaling pathways (**Fig. 2c**). In addition, GO analysis further reveals that downregulated DEGs in acute SIV-infection on short-term ART are related to macrophage activation, Fc-gamma receptor signaling pathway involved in phagocytosis, and positive regulation of macrophage cytokine production, suggesting that short-term ART may alleviate acute SIV-infection induced macrophage activtion responses (**Fig. 2c**). Moreover, toll-like receptor signaling, PI3k/PKB signaling, and osteoclast differentiation signaling are also downregulated (**Fig. 2c**). Of note, no significant DEGs of interest were noticed in the SIV+Short-term ART vs baseline dataset (**SFig. 1**). Altogether, these results indicate that short-term ART effectively alleviates the elevated host antiviral innate immune response, macrophage activation, and the dysregulated T cell and B cell activation to baseline level.

### Long-term ART sustains the downregulated antiviral innate gene expression of short-term ART and upregulates platelet activation and coagulation signaling pathways

To further understand the dynamic transcriptomic changes between chronic SIV infection on long-term ART as ocurred in PLWH vs acute SIV infection as occurred in AIDS patients, we further compared chronic SIV-infected on long-term ART to acute SIV-infected monkeys’ data. Volcano plot and GO analysis show similar changes of DEGs and pathways as acute SIV-infection on short-term ART vs SIV-infection, with upregulated B cell proliferation, negative regulation of T cell proliferation, and downregulated IL27 signaling, antiviral innate response, type I/ type II IFN-mediated signaling pathway, and PI3k/PKB signal transduction (**SFig. 2a-c**). As expected, these data indicate that long-term ART continues to show persistently beneficial effects to alleviate SIV-infection-induced host antiviral innate immunity and reverse SIV-infection-induced dysregulation of adaptive response. Additionally, genes (*F5, ITGA1, ITGA2, ITGA3, MMRN1, TREML1,* etc) and pathways related to coagulation and platelet activation were upregulated, indicating persistent platelet activation and coagulation signaling that is consistent with residual atherothrombotic risk (**SFig. 2a-b**). This is consistent with clinical observation that, compared to healthy individuals, PLWH are more likely to develop CVD, such as atherosclerosis (31), myocardial infarction (48) and stroke (1), with elevated platelet activation markers (49–51), and increased endothelial inflammatory markers (52).

### Chronic SIV infection with long-term ART sustains elevated TLR signaling and macrophage activation compared to baseline or acute SIV with short-term ART

To investigate whether long-term ART returns the host innate immune response to the baseline level, we conducted a pairwise comparison of PBMCs from long-term ART treatment vs baseline monkey PBMC. The volcano plot shows that upregulated genes in long-term ART-treated monkeys include type I IFN signaling gene IRF7 and Cytokine Inducible SH2-containing Protein (*CISH*) (**SFig.3a**). Additionally, OSM, a regulator of the production of other cytokines, including IL6, granulocyte-colony stimulating factor, and granulocyte-macrophage colony stimulating factor in endothelial cells are also upregulated compared to baseline level (**SFig. 3a**). GO analysis further reveals upregulated pathways related to JAK/STAT, NF-kappa B signaling, and PI3k/PKB signaling pathways, and downregulated ribosomal assembly/protein translation related pathways (**SFig. 3b-c**). This suggests that, unlike short-term ART, long-term ART is associated with continuous activation of the immune responses compared to baseline.

**Figure 3.**
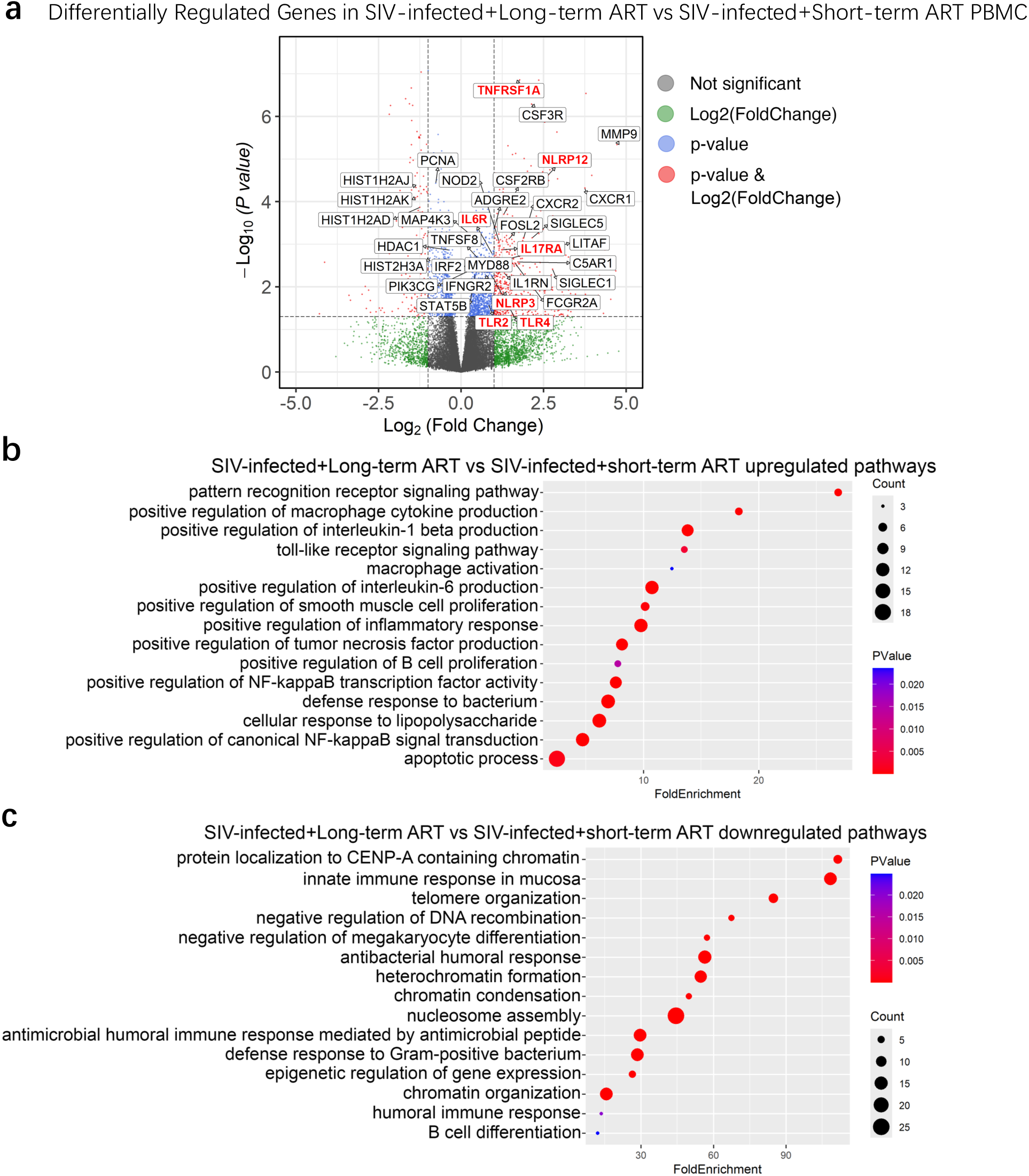
Long-term ART is associated with persistent macrophage activation linked to SIV pathogenesis. **(a)** Volcano plot showing differentially expressed genes (DEGs) in PBMCs of SIV-infected + long-term ART vs SIV-infected + short-term ART. Gene Ontology analysis of **(b)** upregulated and **(c)** downregulated DEGs in long-term ART compared to short-term ART in SIV-infected macaques.

To further understand the long-term effects of ART treatment in SIV-infected individuals, we compared the bulk RNA-seq dataset of chronic SIV infection on long-term ART treatment vs. acute SIV-infection on short-term ART treatment animals. Upregulated genes in chronic SIV infection on long-term ART animals includes genes related to inflammasomes (NLRP3 and NLRP12), colony-stimulating factors (CSF3R, CSF2RB, FOSL2), macrophage activation-related genes (SIGLEC1, SIGLEC5), TLRs and downstream genes (TLR2, TLR4, MYD88), interleukin/interleukin receptor related genes (IL6R, IL17RA), IFN related genes (IFNGR2), LPS induced TNF signaling/TNF receptor related genes (LITAF, TNFRSF1A, TNFRSF8), matrix metalloproteinase (MMP9), and CXC chemokine receptors (CXCR1, CXCR2) (**Fig. 3a**). Downregulated genes are associated with histones modifications (HIST1HAJ, HIST1HAK, HIST1H2AD, HDAC1, HIST2H3A) (**Fig.3a**). Functional analysis further reveals pathway changes in SIV-infected+Long-term ART. Upregulated pathways are related to cellular response to lipopolysaccharide and defense response to bacteria (**Fig. 3a**). Toll-like receptor signaling, especially TLR2/TLR4/MYD88, is also upregulated, as well as downstream signaling pathways such as NF-kB signaling, and further downstream signaling pathways of NF-kB signaling pathways, including IL6 signaling, tumor necrosis factor production, and IL1-beta production pathways, in long-term ART compared to short-term ART (**Fig. 3b**). In addition, NF-KB signaling activated macrophage activation signaling pathway, which is also upregulated in long-term ART compared to short-term ART (**Fig. 3b**). In parallel, the observed downregulated pathways and genes of histone-related genes suggest reduced heterochromatin condensation and epigenomic instability (**Fig. 3c**). This may be related to HIV persistence and latency, leading to persistent protein-driven inflammation. Altogether, this indicates that despite viral suppression with long-term ART, chronic inflammation, persistent microbial translocation, and TLR-driven innate immune activation remain prominent. These changes are consistent with what our and other previous studies have shown, that SIV-infected macaques under long-term ART have persistent inflammation and TLR-driven innate immunity against chronic SIV infection, and ongoing gut disorder (26, 53).

### Persistent atherosclerosis-linked macrophage activation and cell senescence activation are shared features of both acute SIV infection and chronic SIV with long-term ART

To further investigate the changes in macrophage activation and senescence-related signaling at the individual gene level across all phases from baseline till chronic SIV on long-term ART, we created comprehensive heatmaps for the expression level of genes from various macrophage activation signaling-related pathways using MsigDB (v25.1.1)(54, 55). Overall, genes in macrophage activation signaling pathways show increased expression in both acute SIV-infection (Group_2_Pre_ART) and chronic SIV infection with long-term ART (Group 4 LT-ART), but not in pre-infection baseline (Group_1_Baseline), and acute SIV infection with short-term ART (Group_3_ST_ART) (**Fig. 4a**). Specifically, acute SIV-infection and chronic-SIV infection with Long-term ART have comparably more consistently upregulated genes from IFNG/STAT1, and TLR/NF-KB signaling pathways compared to baseline or acute SIV-infection on short-term ART (**Fig. 4a**). Other pathways including antigen processing, type I IFN, IL4/IL13/STAT6, Fc-gamma phagocytosis, complement pathway, NLRP3/IL-1B and IL18 inflammasome signaling pathways still show a similar trend, but with more discrepencies within the biological replicates in the same group (**Fig. 4a**).

**Figure 4.**
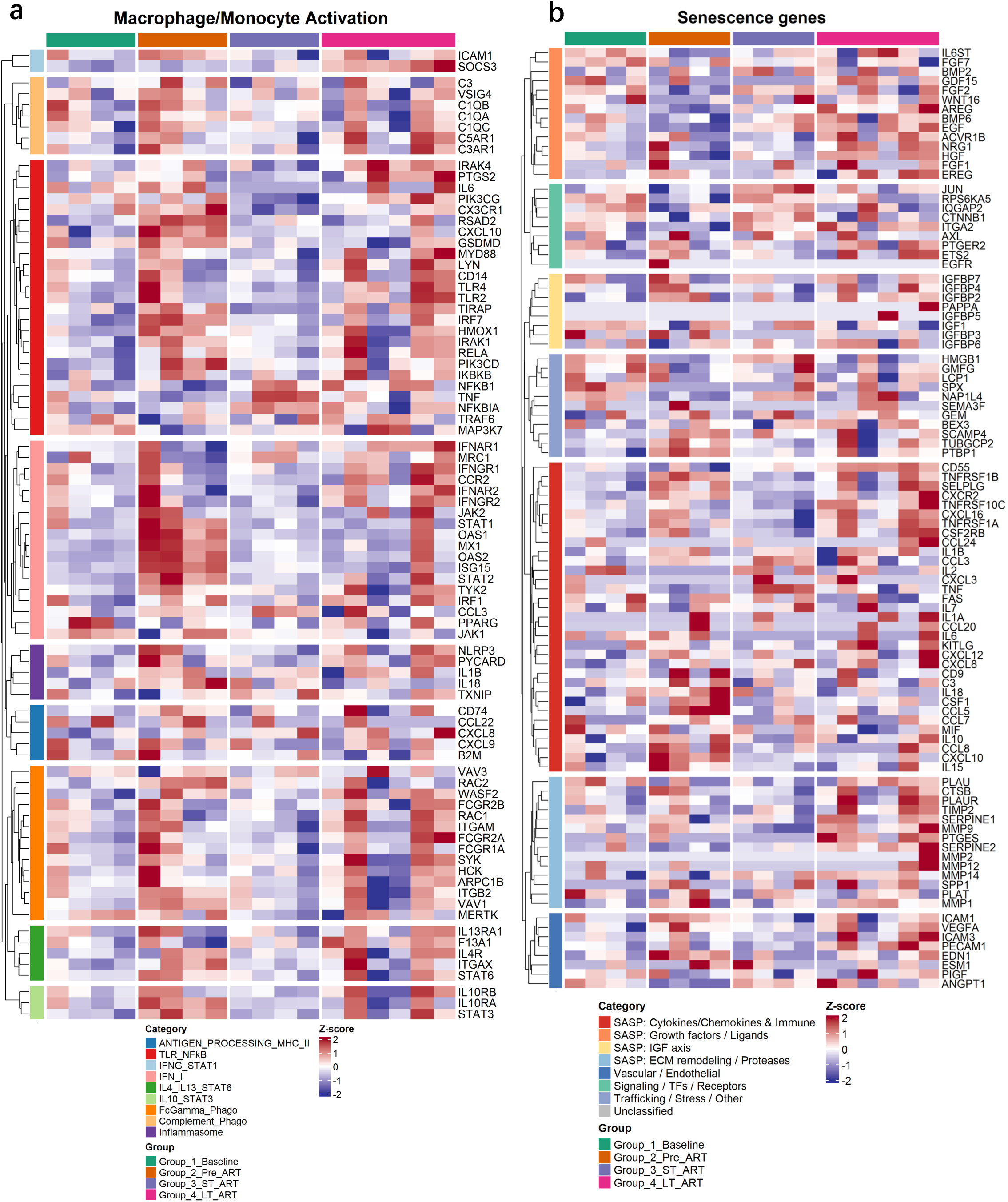
Persistent macrophage activation and senescence programs across the SIV disease course and long-term ART. **(a)** Heatmap of macrophage/monocyte-activation genes mapped to MSigDB signatures (categories include Antigen processing/MHC-II, IFN/NF-κB, IL-1, IL-4/IL-13–STAT6, Fc/complement/phagocytosis,and inflammasome). Z-scores and clustering as in (a). **(b)** Heatmap of curated senescence/SASP genes in PBMCs across four states—Baseline, acute Infection (Pre-ART), Short-term ART, and Long-term ART. Values are per-gene Z-scores of normalized expressions; rows and samples are hierarchically clustered. Left color bars annotate functional categories (e.g., SASP cytokines/chemokines & immune mediators, growth factors/ligands, ECM remodeling/proteases, cell-cycle/DDR, chromatin/TF/receptor modules). Increased expression of multiple SASP modules appears during acute infection and remains detectable in Long-term ART.

**PLWH** are susceptible to inflammatory comorbidities despite effective **ART** and experience premature onset of aging-related comorbidities such as **CVD**. They are twice as likely to develop CVD (1) including atherosclerosis, a leading cause of morbidity and mortality in PLWH. We and others have demonstrated that HIV-induced chronic inflammation contributes to the accelerated aging process and CVD in PLWH (4, 5, 20).

To better understand the mechanism underlying chronic HIV-infection induced atherosclerosis, we further created a heatmap to show the trends in atherosclerosis-associated macrophage signature genes of three conserved major macrophage subsets (Resident/Resident-like, Inflammatory VCAN/IL1B , and Foamy/Trem2hi, IFNIC), which were characterized by a recent single-cell RNA-seq study of the integrated 12 mouse and 11 human patients datasets (56). Heatmap analysis shows an upregulated trend of gene expression such as NLRP3/IL1B in all three subsets of macrophage signatures in acute SIV-infection and in chronic SIV under long-term ART, indicating possible activation of atherosclerosis-related macrophages in those samples (**SFig. 4**).

On the other hand, growing evidence has suggested a premature aging phenotype in PLWH in relation to either HIV disease pathogenesis(10, 57) or highly active antiretroviral treatment (HAART) itself (58, 59) that leads to HIV-associated comorbidities, including atherosclerosis and lung diseases. Heatmap of the SenMayo curated gene list (60) highlights differential expression of senescence-associated secretory phenotype (SASP) genes across groups (**Fig. 4b**). SASP modules related to cytokines/chemokines and immune mediators, IGF-axis ligands, ECM remodeling/proteases, and vascular/endothelial programs, show increased expression in both acute SIV infection and chronic SIV infection with long-term ART (**Fig. 4b**). Consistently, senescence genes related to cell proliferation such as (CTNNB1 etc.) are downregulated in both acute SIV infection and chronic SIV infection with long-term ART (**Fig. 4b**). These patterns indicate a shift toward a pro-inflammatory, pro-remodeling SASP under acute infection and after long-term ART, consistent with persistent innate activation and potential contributions to HIV-related aging process(61).

### Linking chronic inflammation, TLR/NF-κB signaling, macrophage activation and SASP to atherogenesis during chronic SIV infection with long-term-ART

Integrating multilevel analyses across cohorts (**Fig. 5**), PCA separates samples by disease/treatment state with acute SIV infection on short-term ART (ST-ART) shifting toward baseline. In contrast, chronic SIV on long-term ART (LT-ART) is dispersed all over, with three samples remaining distinct and three samples overlapping with short-term ART and baseline (Fig. 5a). This is consistent with previous analysis, showing that short-term ART alleviates host innate immune response and restores B cell and T cell activation to baseline level. At the same time, the LT-ART group partially alleviates host antiviral innate immune response, but has persistent inflammation. Of note, there is a large distribution in the PCA plot between LT-ART samples, which may be related to variation in ART efficacy between different individuals (**Fig. 5a**).

**Figure 5.**
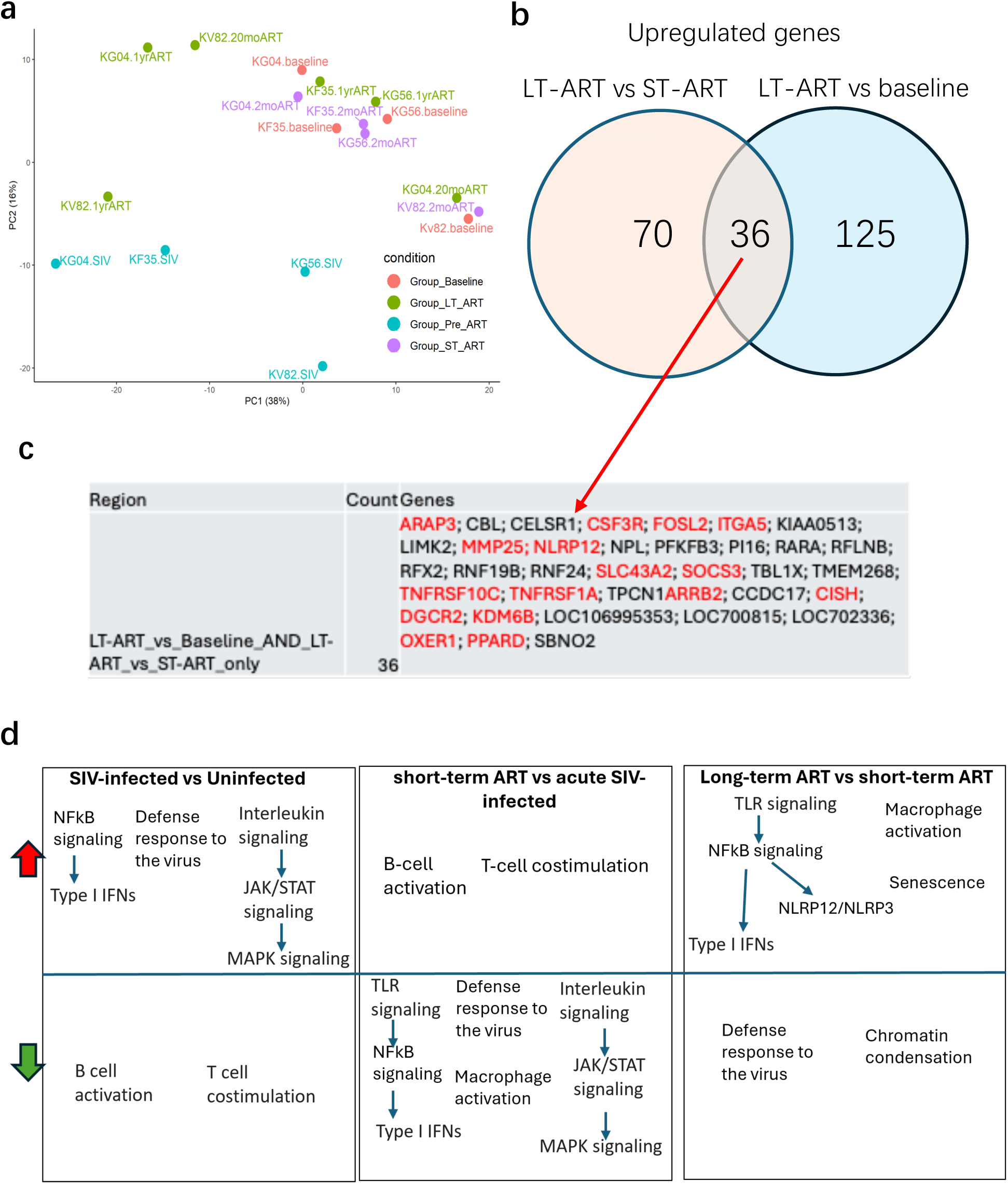
Gene-level overlaps and pathway shifts across acute SIV and ART. **(a)** Principal component analysis of normalized gene expression from four conditions: Baseline (red), acute SIV pre-ART (cyan), short-term ART (magenta), and long-term ART (green). **(b)** Venn diagram of genes significantly upregulated in each comparison: Pre-ART (acute infection) vs Baseline, Long-term ART (LT-ART) vs Baseline, and LT-ART vs Short-term ART (ST-ART). Numbers indicate unique and shared DE genes. **(c)** Genes from key overlap regions. The largest LT-ART–specific set shared with LT-ART vs Baseline and LT-ART vs ST-ART includes macrophage/innate and signaling genes (e.g., **ARAP3, CSF3R, MMP25, NLRP12, SLC43A2, TNFRSF10C, TNFRSF1A**). Additional rows list genes shared by Pre-ART and LT-ART vs ST-ART only, and by all three comparisons. **(d)** Pathway enrichment summaries (GO BP terms) for each contrast. (Pre-ART = acute infection before ART; ST-ART = short-term ART; LT-ART = long-term ART.)

Differential-expression overlaps highlight a recurrent LT-ART signature shared with both LT-ART vs baseline and LT-ART vs ST-ART, enriched for innate/macrophage genes including CSF3R, NLRP12, TNFRSF1A/TNFRSF10C, MMP25, SLC43A2, and FOSL2/ITGA5 (**Fig. 5b–c**). GO pathway summaries show that acute SIV upregulates TLR/NF-κB/type-I IFN programs and dampens B-/T-cell costimulation; ST-ART reverses many of these innate pathways; and LT-ART re-elevates TLR/NF-κB/macrophage-activation and senescence relative to ST-ART (**Fig. 5d**). Together, these alterations point toward a feed-forward inflammatory loop: chronic inflammation, TLR activation, NF-κB signaling, macrophage activation, and SASP cytokine production, sustained systemic inflammation. Functionally, this molecular pattern provides a plausible mechanistic basis for the elevated risk of non-AIDS comorbidities (e.g., CVD, atherosclerosis, aging process) observed in HIV/SIV infection despite prolonged ART.

### Increased activation of caspase-1 and NF-κB in the spleens collected from SIV-infected rhesus macaques on long-term ART

Based on the marked upregulation of NF-κB signaling pathway genes and our previous observation of elevated plasma IL18 and IL1β levels in chronically SIV-infected macaques receiving long-term ART (26), we next investigated whether CASPASE-1 (CASP-1) and NF-κB proteins are activated in the tissues from SIV-infected animals on LT-ART. Western analysis of pro-caspase-1, actived CASP-1, total NF-κB, p-NF-κB were performed on spleen cell lysates from the SIV-infected animals on long-term ART and compared with SIV-naive animals (**Figure 6**). Spleens from SIV+LT-ART exhibited significantly higher levels of active CASP-1 and phosphorylated NF-κB (**Figure 6A**), resulting in an increased CASPASE-1 p20/PROCASPASE-1 and phospho–NF-κB/total NF-κB ratio ratio (**Figure 6B**). Together with the PBMC transcriptomics data described above, these findings demonstrate persistent activation of CASP-1 and NF-κB pathways during chronic SIV infection despite long-term ART, implicating sustained inflammasome and NF-κB signaling, monocyte and macrophage activation, chronic inflammation, and the pathogenesis of HIV-associated CVD and atherosclerosis.

**Figure 6.**
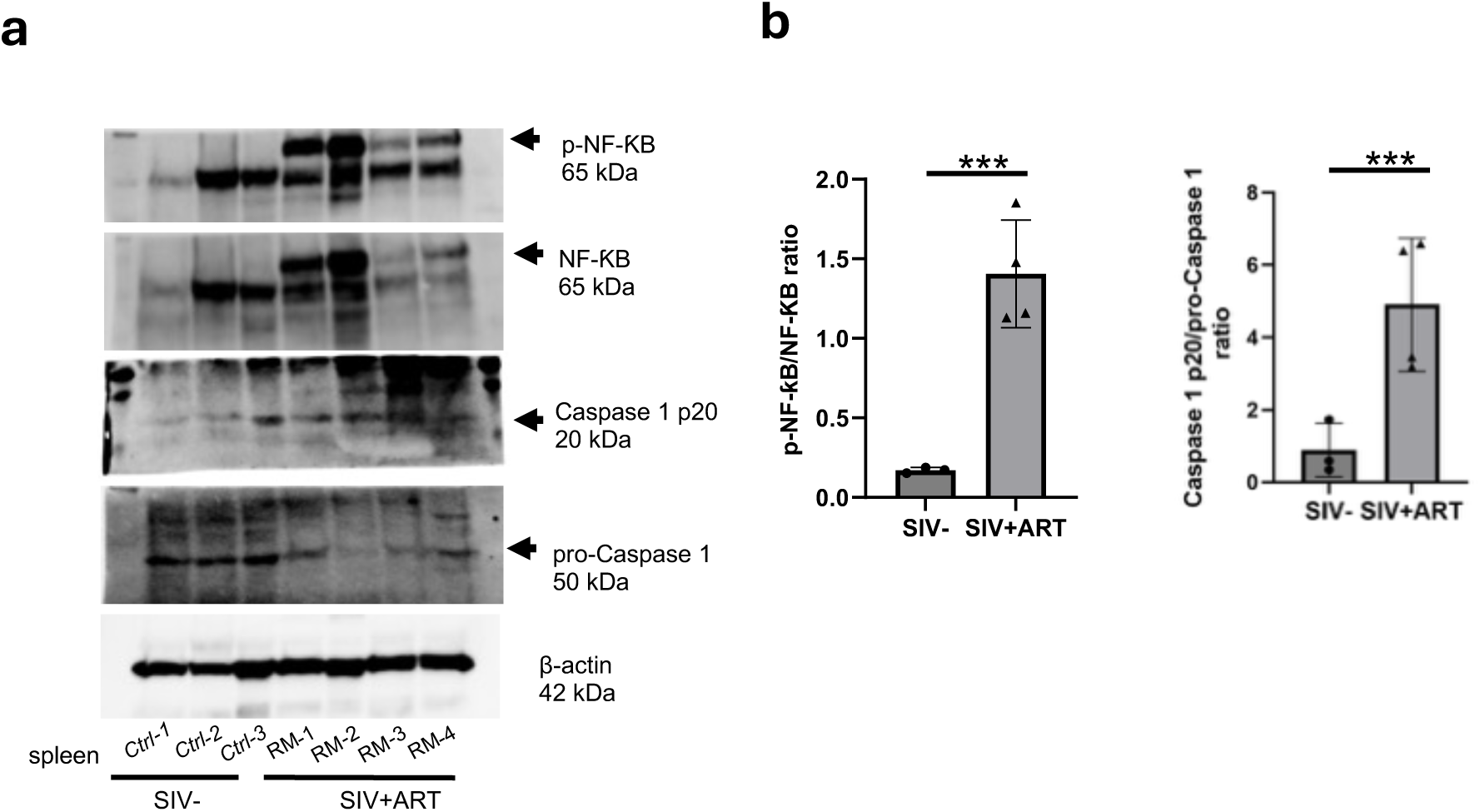
NF-κB and Caspase-1 activation persists in spleens of SIV-infected RMs despite ART. **(a)** Representative Western blot analysis of phosphorylated NF-κB p65 (Ser536), total NF-κB p65, cleaved caspase-1 p20 and procaspase-1 in splenic lysates from uninfected control and SIV-infected RMs on ART. **(b)** Densitometric quantification of the phospho–NF-κB/total NF-κB and caspase-1 p20/procaspase-1 ratio is indicated. Data represent mean ± SEM and P < 0.001 by unpaired t test.

### Cardiovascular pathway activation and macrophage accumulation and senescence in carotid plaque under long-term ART

Canonical Pathway analysis in our cohort further revealed that cardiovascular-related signaling was among the most persistently altered during long-term ART. While early infection and short-term ART showed minimal perturbation, long-term ART induced pronounced activation of adrenomedullin signaling, cardiac β-adrenergic signaling, and platelet homeostasis pathways—key regulators of vascular tone, endothelial integrity, and hemodynamic stress (Supplemental **Fig. 5**). Additionally, pathways involved in cell surface interactions at the vascular wall and factors promoting cardiogenesis were upregulated, suggesting ongoing endothelial activation and tissue remodeling. Furthermore, stathmin1-associated and cancer-related signaling modules suggested the emergence of proliferative and senescence-linked transcriptional programs. Overall, these data point to sustained vascular and cardiac stress responses under long-term ART, consistent with chronic inflammation and potential predisposition to cardiovascular dysfunction despite effective viral suppression.

To establish the clinical relevance of our transcriptomic findings, specifically the activation of macrophage accumulation, senescence, platelets and coagulation, and NLRP3-CASAPSE-1 pathways in the context of HIV-associated CVD, we performed a pathological characterization of a carotid atherosclerotic plaque from an SIV-infected male rhesus macaque (7 year old) on a similar LT-ART regimen for 7 months. We observed small atherosclerotic plaques (fatty streaks, potentially type II by AHA classification(62)) (**Figures 7A** and **7B1**). Plaque composition analysis showed key features aligning with our transcriptomics results including the presence of abundant F4/80 and CD68-immunopositive macrophages (**Figure 7B2**), p21-positive senescent cells (**Figure 7B3**), and low collagen and αSMA-positive smooth muscle cells (SMC) (**Figure 7B3**), as well as an CD31-positive endothelial layer with multiple breaks (**Figure 7B3**). In addition, the immunopositivity for IL-1 beta, a product of NLRP3-CASPASE 1 activation was strongly associated with vascular SMC’s marker positivity in carotid media and with cells in the fibrous cap in the plaque (**S. Fig. 6**). In the carotid, NLRP3+ signal was strongly associated with cells in the plaque (**S. Fig. 7**). Surprisingly, we also found a thrombus within the artery (**Figure 7A** and **7C**). This thrombus contained collagen (**Figure 7C1**), micro-vessels (**Figure 7C2, right image**) that was immunopositive for thrombin, a key blood clot-forming enzyme (**Figure 7C2, left image**), and RBCs (**Figure 7C**). Given the small sized plaque, rupture is an unlikely cause for such a large thrombus. Nevertheless, this unexpected observation in the index SIV-infected macaque provides pathological evidence of atherothrombotic disease in carotid artery, along with macrophage accumulation and senescence in a carotid plaque during chronic SIV infection with long-term ART. Overall, these pathological findings align with our transcriptomic profile of heightened monocyte/macrophage, senescence, and platelet/coagulation pathway activation in SIV-infected animals on long-term ART.

**Figure 7.**
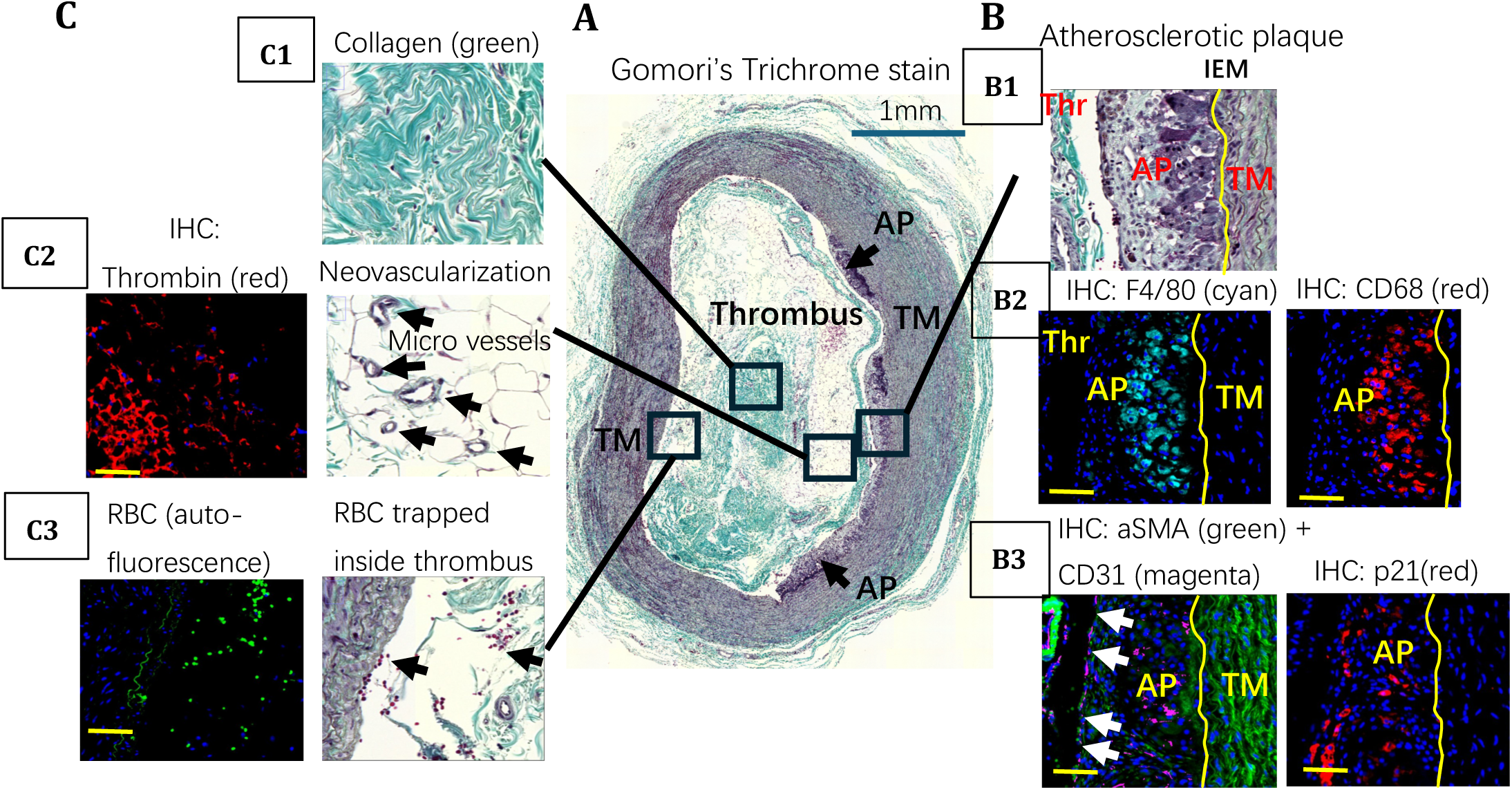
Vascular morphology and atherosclerotic plaque (AP) composition in carotid artery of macaque rhesus under chronic SIV+ART treatment. A, Trichrome staining detects the presence of collagen and neovascularization in occlusive thrombus (Thr) and atherosclerotic plaques (AP) (black arrows) developed on the tunica media (TM); **B,** Immunohistochemistry (IHC) was performed using antibodies against F4/80 and CD68 (macrophage markers) **(B2)**, aSMA (SMC marker) and CD31 (endothelial cell marker) **(B3)** and p21 (senescent cell marker). IHC shows the colocalization of F4/80-positive and CD68-positive macrophages with senescent p21+ cells in AP. aSMA+ cells were detected mainly in TM and CD31+ endothelial layer contains numerical cells breaks (white arrows) **(B3)**. Yellow curve outlines internal elastic membrane (IEM), the borderline between TM and AP; **C,** the presence of occlusive thrombus was confirmed by IHC using antibody against thrombin **(C2)**. RBC trapped inside thrombus were visualized by autofluorescence with FITC filter. Yellow scale bar, 50um.

## Discussion

Despite effective viral suppression with ART, PLWH experience persistent inflammation and immune dysfunction that drive long-term comorbidities, underscoring the importance of elucidating immune trajectories during infection and treatment. In this study, we provide a comprehensive transcriptomic and molecular characterization of immune responses across the full course of SIV infection, from acute infection through short- and long-term ART, in a longitudinal nonhuman primate model. By integrating bulk RNA-seq of PBMCs, pathway enrichment analyses, and validation at the protein and tissue levels, we delineate how SIV infection induces dynamic, phase-specific immune remodeling. Our findings demonstrate that acute SIV infection triggers robust antiviral innate immune activation and interferon responses including ISGs, IL27 signaling, NF-κB signaling, and proinflammatory chemokines, while downregulating T- and B-cell activation pathways, indicating suppression of adaptive immunity. While these responses are progressively normalized by ART, persistently altered innate and inflammatory pathways, including TLRs/NF-κB, inflammasome, macrophage activation, and senescence programs, remain active during long-term ART, paralleling the chronic inflammation and age-related comorbidities observed in PLWH.

Consistent with prior studies of acute HIV/SIV infection (63, 64), we observed dramatic upregulation of type I and type II ISGs, antiviral response genes (e.g., IFI44, ISG15, DDX58/RIG-I, MAVS), and inflammatory mediators (CXCL10, IL27, STAT1). These changes highlight the dominance of IFN-driven antiviral defense and innate activation during early infection. Concomitantly, genes associated with T-cell and B-cell activation (CD28, CD72) were downregulated, and inhibitory immune checkpoint genes (PDCD1, CD276) were upregulated, indicating suppression of adaptive immunity and early onset of immune exhaustion. This SIV-induced adaptive immune suppression is consistent with the marked and sustained CD4⁺ T-cell depletion characteristic of HIV/SIV infection, which disrupts T helper function essential for B-cell activation and antibody production. GO analysis confirmed these patterns, showing enrichment for interferon, JAK/STAT, NF-κB, and MAPK signaling pathways. Collectively, these results demonstrate that acute SIV infection activates a potent antiviral innate immune program while disrupting T- and B-cell stimulation, a pattern characteristic of early HIV pathogenesis in humans (65, 66). It is important to note that we also demonstrate that IL-27-mediated signaling pathway is the most upregulated pathway after SIV infection (Figure 1E). This upregulation is normalized by either short (most downregulated pathways in SIV-infected+Short-team ART vs SIV-infected PBMC down-regulated pathways shown in Figure 2C) or long-term ART treatment (highest down-regulated pathways in chronic SIV-infection on long-term vs SIV-infected PBMC down-regulated pathways). This result clearly indicates that IL27-mediated signaling pathway activation is the most important host response to the initial SIV or HIV infection. Consistently, previous studies have demonstrated that the IL27 pathway has inhibitory and promoting effects for HIV infection in PBMCs (67, 68) and it may have a theraputic potential for treating HIV infection (69, 70). Therefore, these results highlight the importance of IL-27-mediated signaling pathway activation in HIV-infection. Its exact role and therapeutic potential in HIV-infection warrants further investigation.

Consistent with several other reports, with the effective suppression of viremia following ART, many of the transcriptional alterations induced by SIV infection were reversed in our cohort (71, 72). Genes related to B-cell activation (CD79A/B, CD40, CD72) and T-cell costimulation (CD28, ICOS, DPP4) were upregulated, while inhibitory checkpoint (PDCD1, CD276) and ISGs were downregulated. These shifts were accompanied by suppression of inflammatory pathways including type I IFN, NF-κB, and IL6 signaling, along with reduced macrophage activation and Fc receptor–mediated phagocytic pathways. Functionally, these results demonstrate that ART rapidly reestablishes immune homeostasis by dampening hyperactive innate responses and restoring adaptive immune activation, consistent with clinical observations of immune reconstitution following ART initiation (71, 72). These findings not only validate the immunodynamic parallels between SIV-infected macaques and HIV-infected humans but also establish a foundation to explore the long-term immunologic consequences of sustained viral suppression. To our knowledge, the progressive, longitudinal transcriptomic remodeling that occurs within individuals during long-term ART-suppressed SIV or HIV infection has not been systematically characterized.

Although long-term ART effectively maintained viral suppression and durable downregulation of ISGs, our transcriptomic and pathway analyses revealed persistent activation of inflammatory, coagulation, and macrophage-associated programs. Notably, genes involved in platelet activation and coagulation (F5, ITGA2, TREML1, MMRN1), innate immune sensing (TLR2, TLR4, MYD88), inflammasome components (NLRP3, NLRP12), and macrophage activation (SIGLEC1, CSF3R) were upregulated. Pathway enrichment analyses further highlighted sustained activation of Toll-like receptor, NF-κB, and TNF signaling, indicating ongoing TLR-driven inflammation and macrophage activation despite prolonged ART (73–75). This chronic innate immune activation closely mirrors the clinical phenotype of PLWH, who exhibit persistent systemic inflammation, endothelial dysfunction, and elevated risk of atherosclerosis, myocardial infarction, and stroke (5, 11). In this study, we have functionally validated the increased CASPASE-1 and NF-κB activation in spleen tissue of chronic SIV on long-term ART treatment, confirming inflammasome and inflammatory pathway persistence under long-term ART. This study extends our previous observation showing the increased CASPASE-1 and NF-κB activation in spleen tissue of acute SIV on short-term ART treatment (20, 21). These results suggest that viral suppression alone does not fully restore immune function (76). Consistently, increased levels of CASPASE-1 and IL-1β in the circulating immune cells has been reported in PLWH with persistent immune activation even after 12 years of suppressive ART (77). Besides, our prior studies have demonstrated increased levels of plasma IL-1β and IL-18 during long-term SIV+ART (26) as well as correlation of CASPASE-1^+^ cells and monocyte/macrophage activation in HIV-patient-derived aortic plaques (20). These data support a model in which long-term ART preserves viral control but maintains low-grade inflammation through macrophage and platelet activation, promoting tissue remodeling and vascular injury, likely drived by persistent viral infection and latency in the tissues (78–80). Our findings align with prior evidence that chronic HIV infection promotes monocyte and macrophage activation via microbial translocation, TLR stimulation, and inflammasome engagement (81–83). The upregulation of NLRP3 and IL1B observed here, together with sustained NF-κB signaling, supports a feed-forward inflammatory loop. This mechanism likely contributes to the senescence-associated secretory phenotype (SASP) observed in both acute and chronic SIV infection, consistent with premature aging and inflammatory comorbidities in ART-treated PLWH. Indeed, the transcriptomic signatures of long-term ART included increased expression of senescence and atherosclerosis-associated macrophage markers, paralleling clinical observations of heightened cardiovascular and thrombotic risk (3, 5, 11, 20, 84).

Histologic analysis of carotid lesions in an index SIV-infected macaque on long-term ART revealed macrophage-rich plaques with senescent cells. Furthermore the presence of intraluminal thrombus suggests a pro-inflammatory and pro-thrombotic state. These findings provide direct evidence linking persistent macrophage activation, SASP, and vascular pathology in the context of ART-suppressed SIV infection. Together, the integration of transcriptomic, proteomic, and histopathological data underscores a central concept: long-term ART resolves viral replication but does not restore immune homeostasis, instead maintaining a chronic inflammatory state that promotes vascular injury and immune aging (3, 5, 11, 20). Previous studies have identified that the accumulation of macrophage foam cells in the intima is instrumental to lesion development, and inflammatory macrophages are crucial for myocardial infarction development, and resident/resident-like macrophages are associated with carotid endarterectomy plaques (56, 85–87). It is important to note that young rhesus macaques fed a standard chow diet rarely develop atherosclerotic lesions or intraluminal thrombi in the carotid artery (88–91). Although, the observed thrombus may not be directly attributable to the relatively small plaque identified in this study animal with chronic SIV+ART, an increased risk of thrombosis has been reported in chronic HIV infection on ART (2) (3). Further, enhanced platelet and coagulation pathway activation and monocyte and macrophage activation significantly contribute to atherosclerosis-associated CVD, including acute coronary attack and stroke (1, 4). Our findings of macrophage accumulation, IL-1beta staining in fibrous caps of the plaques, NLRP3 nuclear location in the plaques cells, and p21⁺ senescent cells with intraluminal thrombus formation in the carotid artery of an ART-suppressed SIV-infected macaque recapitulate key features of HIV-associated atherogenesis and thrombosis. However, the cellular and molecular mechanisms (eg: NLRP3-CASPASE-1, NF-kb, senecence, platelet and coagulation pathways) driving HIV-associated atherosclerosis and thrombosis remain incompletely defined and warrant further study in clinically relevant nonhuman primate models (5). To enable more systematic investigation of these processes within an experimentally tractable timeframe, a refined model employing ART-suppressed, chronically SIV-infected rhesus macaques fed an atherogenic diet (85) (92) may provide a valuable platform for dissecting the pathogenic mechanisms of HIV-associated thrombosis and atherosclerosis.

In summary, this study provides comprehensive longitudinal evidence from a preclinical model of HIV infection, delineating distinct immunologic phases from acute infection through chronic ART-suppressed disease. We identify persistent Caspase-1 activation driven by NLRP3–NLRR12 inflammasome formation, together with NF-κB signaling, monocyte/macrophage activation, cellular senescence, and platelet/thrombosis pathway engagement, as hallmarks of chronic immune dysregulation under ART. These processes likely contribute to the heightened cardiovascular, thrombotic, and immunosenescence-related comorbidity risk observed in PLWH. Collectively, our findings highlight TLR–NF-κB signaling, inflammasome activity, and senescence pathways as potential therapeutic targets to mitigate chronic inflammation and reduce non-AIDS comorbidities in ART-treated HIV infection.

## Methods

### Sex as a biological variable

Our cohort is a part of previous studies that used only female macaques. However, we have observed similar temporal dynamics of the immune cell subpopulations and gut barrier disruption in male macaques in our other cohorts utilizing identical SIV infection and ART regimen.

### Animals, Viral Inoculation, and ART

Six healthy adult female Indian-origin rhesus macaques (*Macaca mulatta*), aged 5–10 years and seronegative for SIV, HIV-2, STLV-1, SRV-1, and herpes-B, were infected intrarectally with 2500 TCID50 SIVmac251 (Preclinical Research and Development Branch, NIAID). ART was administered daily via subcutaneous injection: 5.1 mg/kg Tenofovir Disoproxil Fumarate (TDF), 30 mg/kg Emtricitabine (FTC), and 2.5 mg/kg Dolutegravir (DTG) in 15% kleptose solution at pH 4.2 (53). Plasma viral loads were quantified using the Roche High Pure Viral RNA Kit (93).

### Cell Isolation

Blood collected in EDTA tubes (Sarstedt) was processed immediately. PBMCs were isolated by density gradient centrifugation (Lymphocyte Separation Medium, MP Biomedicals) at 1500 rpm for 45 min for phenotyping and functional assays. Splenocytes were isolated from tissue obtained at necropsy. Briefly, pieces of tissue were washed with phosphate buffered saline (PBS), minced and homogenized with a syringe plunger and filtered through a cell strainer. The cell suspension in the filtrate was washed and resuspended in RPMI-10 + 10% FCS. Cell viability was >90% by trypan blue exclusion.

### RNA isolation

Tissues were collected in 1 mL Trizol reagent (15596026; Invitrogen) and extracted with RNeasy Mini Kit (Cat. No.74104; QIAGEN, Hilden, Germany) following the manufacturer’s protocol. The concentration of RNA was determined by NanoDrop 2000.

### Bulk RNA-seq analysis

Final cDNA libraries carrying TruSeq RNA CD indexes (Illumina, 20019792) were quantified with the Qubit dsDNA HS Assay Kit (Thermo Fisher Scientific, Q32854). Library quality was assessed on an Agilent TapeStation 4150 using Agilent D1000 ScreenTape (Agilent, 5067-5582). Smear analysis (Agilent TapeStation Software, Version 4.1.1) with a 600 bp window was used to determine the average library size. Library molarity was then calculated from the measured size and concentration. Libraries were pooled to a final concentration of 750pM with a 2% spike-in of PhiX control library v3 (Illumina, FC-110-3001). The pooled mixture was loaded onto an Illumina NextSeq P1 (300) reagent cartridge (Illumina, 20050264). Paired-end, dual-index sequencing (150×8×8×150) was run on the NextSeq2000, generating ∼20 M paired-end reads per sample. Fastq files produced by Illumina BaseSpace DRAGEN Analysis Software (Version 1.2.1) were used for downstream analyses. Raw reads were evaluated with FastQC and aligned to mm10 using Hisat2. Transcript assembly and abundance estimation were performed with the featurecounts R package. Gene-level differential expression was analyzed with DESeq2. Volcano plots were created with the Enhanced Volcano package (v1.16.0) to visualize gene-level changes. Gene ontology (GO) analysis used DAVID Functional Annotation Bioinformatics Microarray Analysis; gene sets with p-value < 0.05 were deemed significantly enriched, and results were visualized with enrichplot, ggplot2, and base R graphics.

### Western blotting

Spleens from untreated or SIV-infected rhesus macaques with ART were homogenized in Pierce RIPA buffer (89900, Thermo Fisher Scientific) supplemented with a protease and phosphatase inhibitor cocktail (5872, Cell Signaling Technology) using a FastPrep-24 5G homogenizer (MP Biomedicals). Total protein concentrations were determined by the Pierce BCA Protein Assay Kit (23227, Thermo Fisher Scientific). Equal amounts of protein were resolved on 4-20% polyacrylamide gels (Bio-Rad) and transferred onto PVDF membranes (Bio-Rad). Membranes were blocked with EveryBlot Blocking Buffer (Bio-Rad), incubated with primary antibodies overnight at 4°C and HRP- or fluorescence-conjugated secondary antibodies for 1 h at room temperature. Signals were developed with the SuperSignal™ West Atto Ultimate Sensitivity Substrate (A38554, Thermo Fisher scientific) and visualized with a ChemiDoc MP Imaging System (Bio-Rad). Primary antibodies were as follows: rabbit mAb against phospho-NF-κB p65 (Ser536) (93H1) (#3033, Cell Signaling Technology), NF-κB p65 (D14E12) XP^®^ (#8242, Cell Signaling Technology) and CASPASE 1 p20 (Cleaved Asp296) (PA5-99390, Invitrogen), and mouse mAb antibody against PROCASPASE-1 (D-3) (sc-392736, Santa Cruz Biotechnology). Secondary antibodies included anti-rabbit IgG, HRP-linked Antibody (#7074, Cell signaling technology), goat anti-mouse IgG StarBright™ Blue 700 (12004158, Bio-Rad), and anti-mouse IgG, HRP-linked antibody (#7076, Cell Signaling Technology).

### Tissue Histology and Immunohistochemistry (IHC)

To visualize collagen, carotid cross-sections were stained with Gomori’s Trichrome stain (cat# 24205, Polysciences). To perform IHC serial carotid artery sections were deparaffinized, dehydrated and processed with heat-mediated antigen retrieval using citrate buffer (pH 6.0) followed by blocking step (Protein block, Abcam, ab64226). Sections were incubated overnight at +4oC with primary antibody or with normal IgG (negative control). The primary antibodies used for IHC are: mouse anti-thrombin antibody (Abcam, clone 5G9, cat# ab17199), rat F4/80 antibody (Abcam, clone CI: A3-1, cat#6640), mouse anti-CD68 antibody (Abcam, clone KP-1, ab955), rabbit anti-p21 antibody (Proteintech, cat#10355-1-AP), rabbit anti-IL-1beta (cleaved Asp116) antibody (Thermo Fisher, cat#PA5-105048), rabbit NLRP3 antibody (Proteintech, cat#19771-1-AP), rabbit anti-CD31 antibody conjugated with AlexaFluor 647 (Abcam, clone EP3095, cat#abd310240), mouse α-smooth muscle actin (αSMA) antibody conjugated with AlexaFluor 488 (Abcam, clone 1A4, cat#ab184675). The primary antibody signal was amplified with horse anti-rabbit-biotin IgG (Vector Laboratories, BP-1100-50) or goat anti-mouse-biotin IgG (Vector Laboratories, BP-9200-50) or goat anti-rat-biotin IgG (Vector laboratories, BA-9400-1.5) followed by incubation with streptavidin-AlexaFluor 594 conjugate (Life Technologies, S32356) plus DAPI. Sections were mounted with ProLong Gold antifade media (Thermo Fisher, P36970) for imaging. The sections were scanned with Cytation 5 multi-mode imager (Bio-Tek, Winooski, MI) using standard TexasRed, GFP and DAPI filter cubes to generate greyscale 16-bit TIFF images for each channel. RBC is known to have strong background signal (autofluorescence) when sections imaged with FITC filter (PMID: 29058770). To visualize RBC, deparaffinized and dehydrated monkey sections were stained with DAPI and imaged with GFP filter.

### Statistical analysis

Densitometric quantification of Western blot bands was performed using ImageJ.JS (NIH), and statistical analyses and data visualization were conducted using GraphPad Prism 10 software (GraphPad Software, San Diego, CA). Data are presented as mean ± SEM. Statistical significance was determined using unpaired two-tailed *t* tests or one-way ANOVA, with p < 0.05 considered significant.

### Study approval

All animal studies were approved by the Tulane University Institutional Animal Care and Use Committee, and all animals were born and housed at the Tulane National Primate Research Center in accordance with the Association for Assessment and Accreditation of Laboratory Animal Care International standards. Animal housing and studies were carried out in strict accordance with the recommendations in the Guide for the Care and Use of Laboratory Animals of the National Institutes of Health (NIH, AAALAC #000594) and with the recommendations of the Weatherall report: The Use of Non-Human Primates in Research.

### Data Availability Statement

Bulk RNA sequencing data reported in this paper will be deposited to the GEO database after the acceptance and will be available publicly at the date of publication (accession number: pending). Relevant information about data available directly upon request to the corresponding authors.

## Author contributions

NR, XQ and JR developed the concept. YC, DX, SR, ST, RB, AS, and SS contributed to perform the experiments and analyzed the results. YC, SS, NR, and XQ wrote the manuscript and all authors participated in the review and critique of the manuscript. NR, SS, PD and XQ interpreted the results and supervised the experiments.

## Acknowledgments

We thank Kejing Song and Haiyan Miller for technical assistance related to Next Generation Sequencing. We also thank confocal microscopy and molecular pathology core: RRID: SCR_024613; Anatomic Pathology Core: RRID: SCR_024606; Virus Characterization, Isolation, Production and Sequencing Core: RRID:SCR 024679; and Confocal Microscopy and Molecular Pathology Core: RRID:SCR 024613; at Tulane National Biomedical Research Center for technical assistance related to histological analysis.

