## Supplemental Files for "Persistent Activation of Monocytes/Macrophages and Cell Senescence in SIV-Infected Macaques on ART"

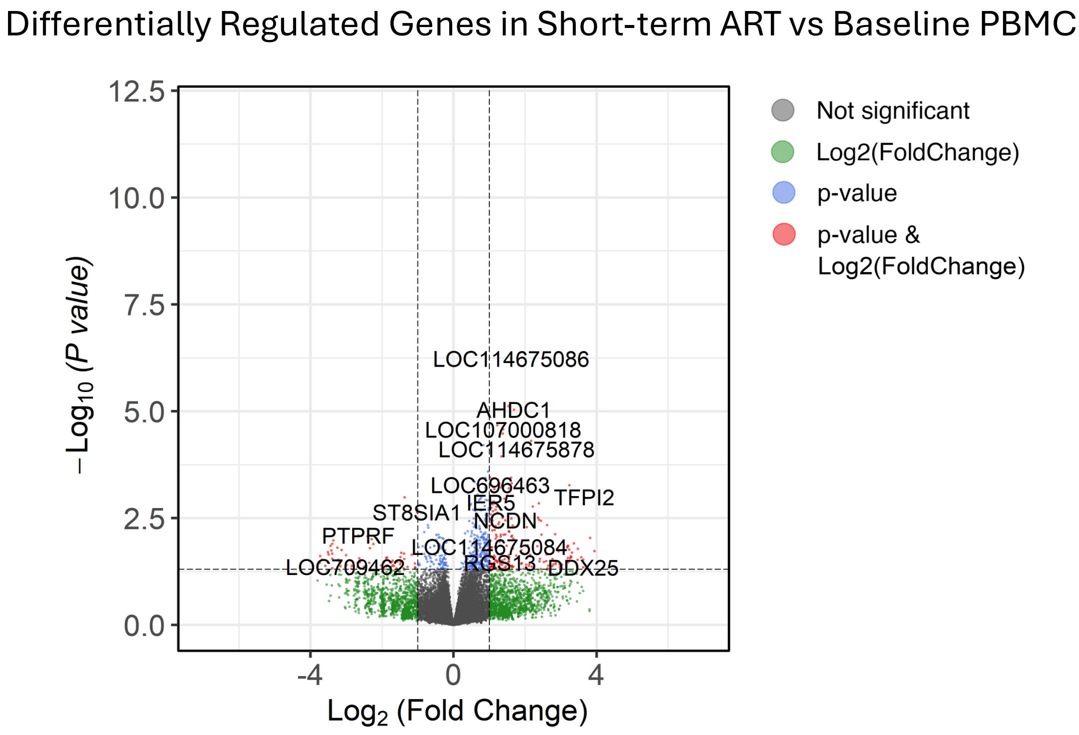


**Supplemental Figure 1. Differential gene expression in PBMCs after short-term ART versus baseline.** Volcano plot showing differentially expressed genes (DEGs) in SIV-infected + short-term ART vs baseline PBMC.

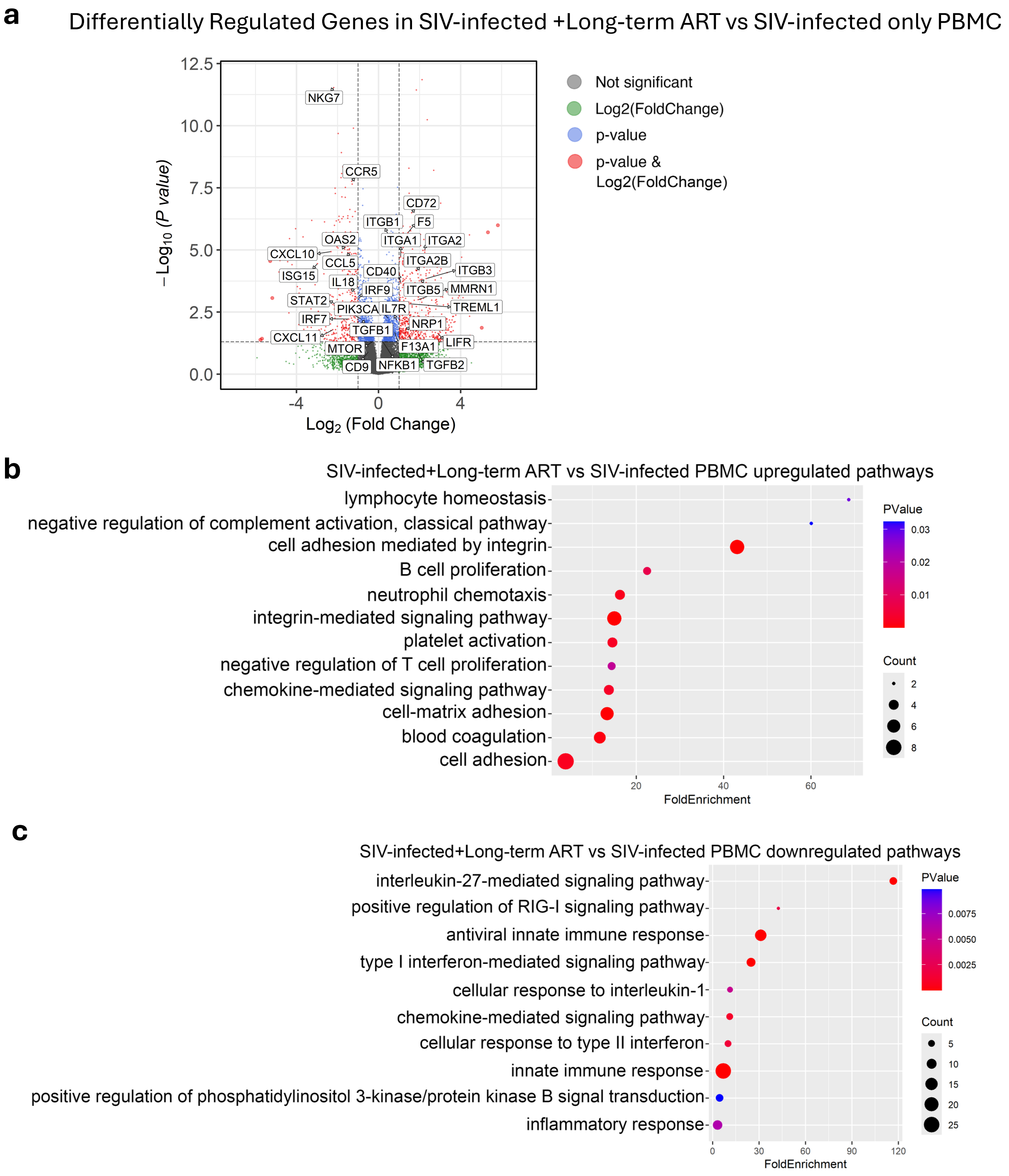


**Supplemental Figure 2. Long-term ART treatment continuously downregulates host antiviral innate immune responses but upregulates integrin and coagulation-related signaling pathways. (a)** Volcano plot showing differentially expressed genes (DEGs) in SIV-infected + long-term ART vs SIV-infected monkey PBMC. **(b-c)** Gene Ontology analysis of **(b)** upregulated and **(c)** downregulated DEGs in SIV-infected + long-term ART monkeys compared to monkeys PBMC.

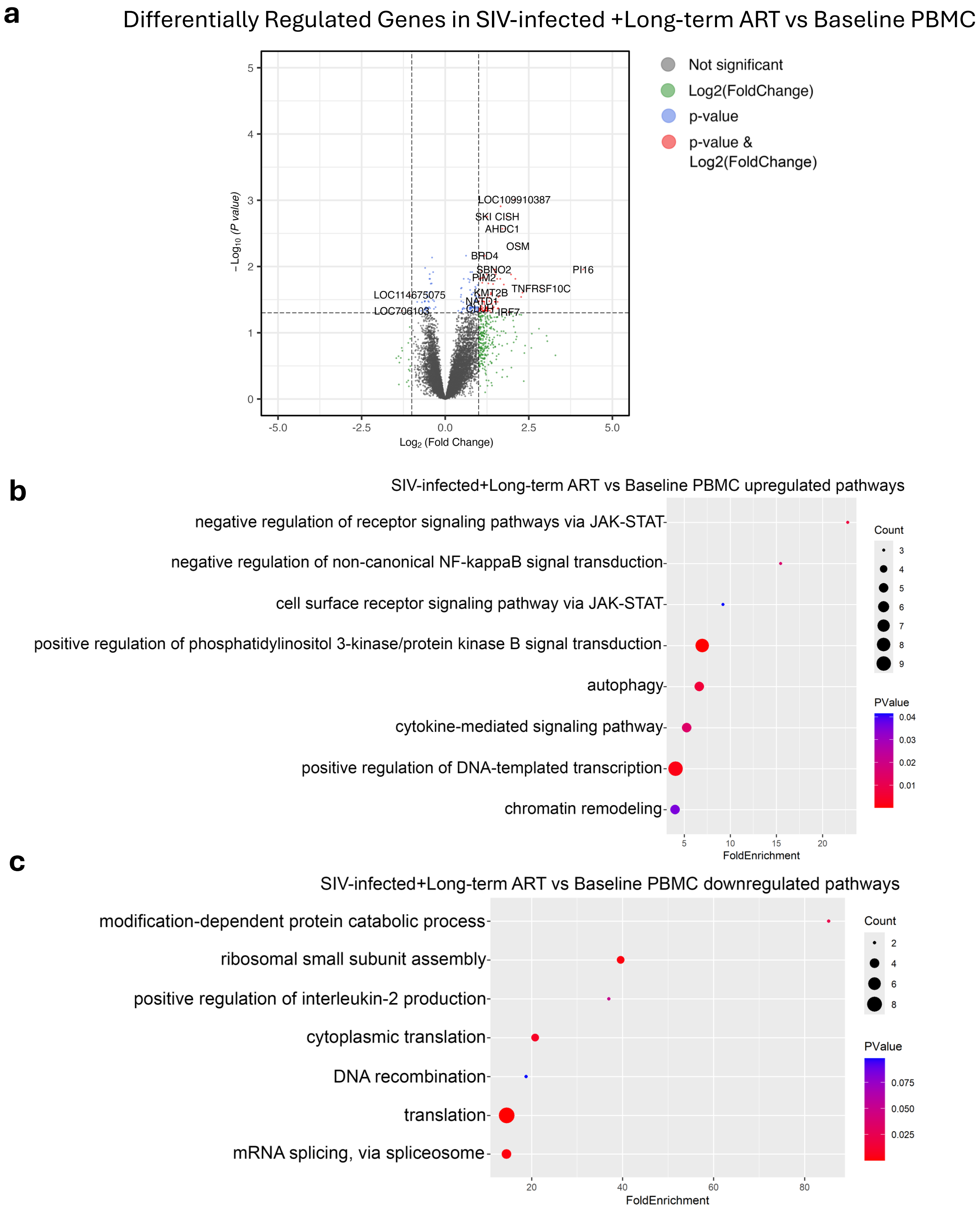


**Supplemental Figure 3. Differential expression and pathway enrichment in PBMCs from SIV-infected animals on long-term ART versus baseline. (a)** Volcano plot showing differentially expressed genes (DEGs) in SIV-infected + long-term ART vs baseline monkey PBMC. **(b-c)** Gene Ontology analysis of **(b)** upregulated and **(c)** downregulated DEGs in SIV-infected + long-term ART monkeys compared to baseline monkeys' PBMC.

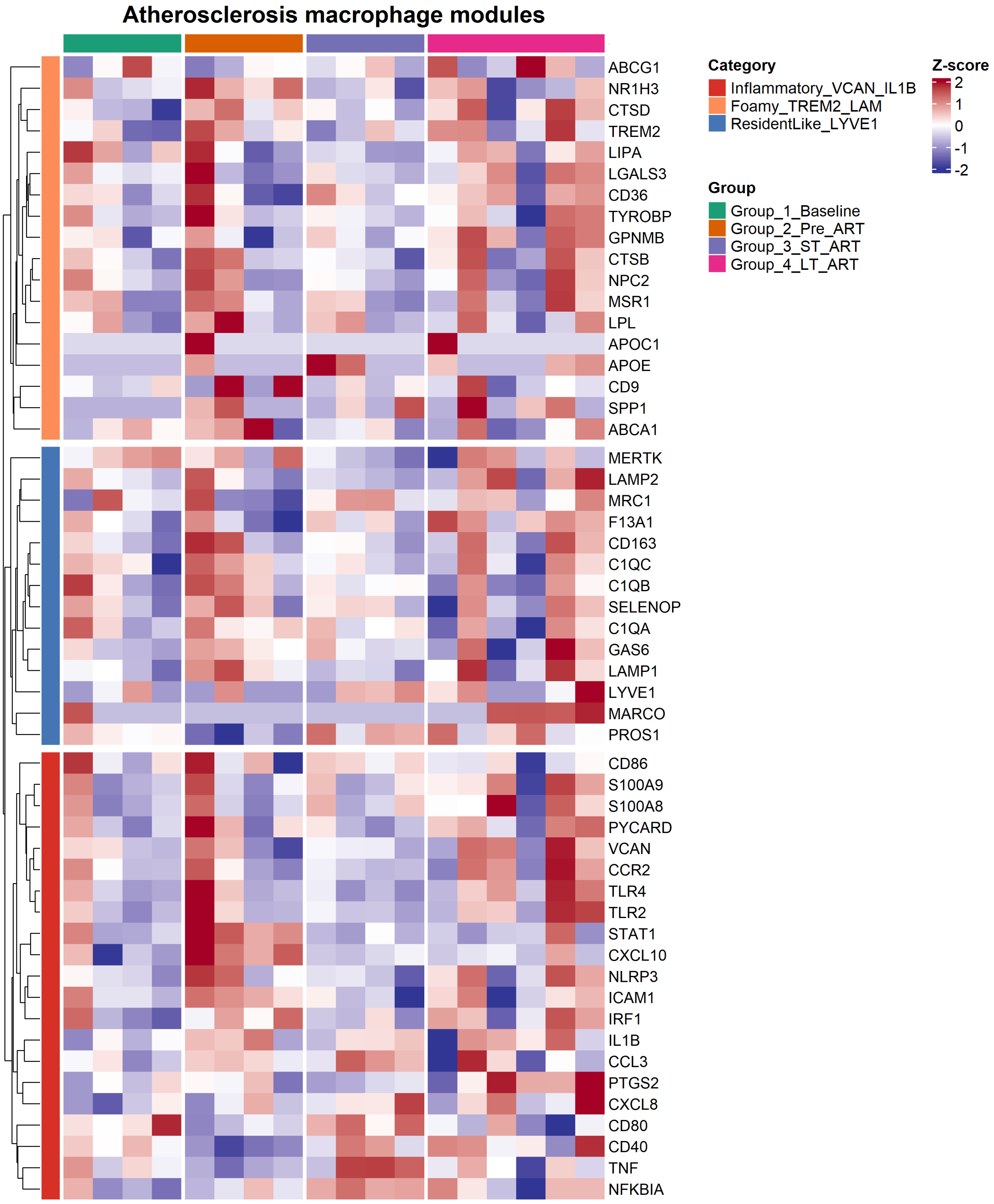


**Supplemental Figure 4. Atherosclerosis-associated macrophage modules across SIV and ART states.** Heatmap of curated atherosclerosis-associated macrophage signature genes grouped into three: Inflammatory_VCAN/IL1B, Foam_TREM2/LIPA, and Resident_LYVE1 (left color bars).

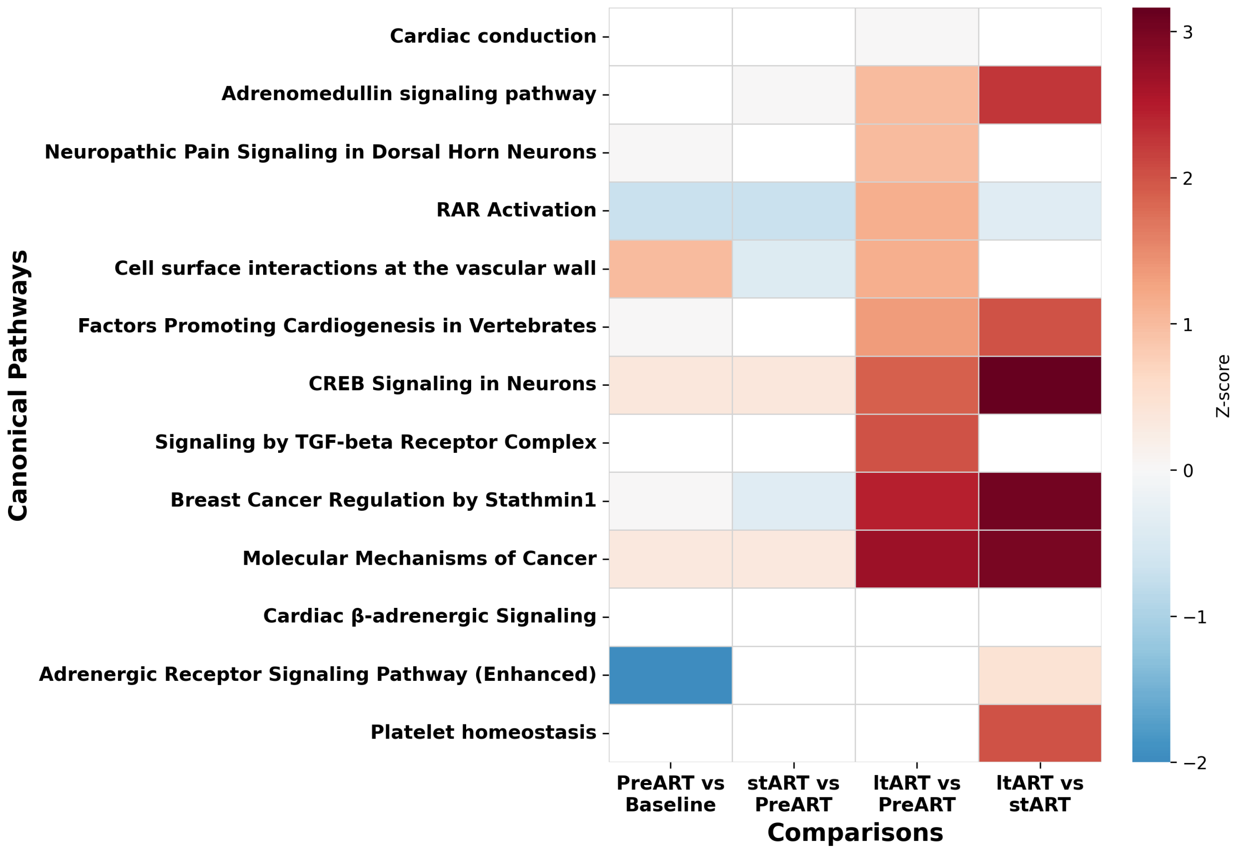


**Supplemental Figure 5. CVD-associated pathways and atherogenesis during Chronic SIV+ART.** Canonical pathway activity across infection and treatment stages showing persistent activation of cardiovascular signaling pathways during ltART. Heatmap shows predicted activation (red) or inhibition (blue) Z-scores of canonical pathways involved in vascular and cardiac regulation across sequential stages: PreART vs Baseline, short-term ART (stART) vs PreART, long-term ART (ltART) vs PreART, and ltART vs stART. While short-term ART induced limited changes, long-term ART was marked by strong activation of adrenomedullin, β-adrenergic, and platelet homeostasis pathways, as well as vascular interaction and cardiogenic signaling programs. These findings indicate persistent vascular and cardiac stress signaling during chronic infection and prolonged ART.

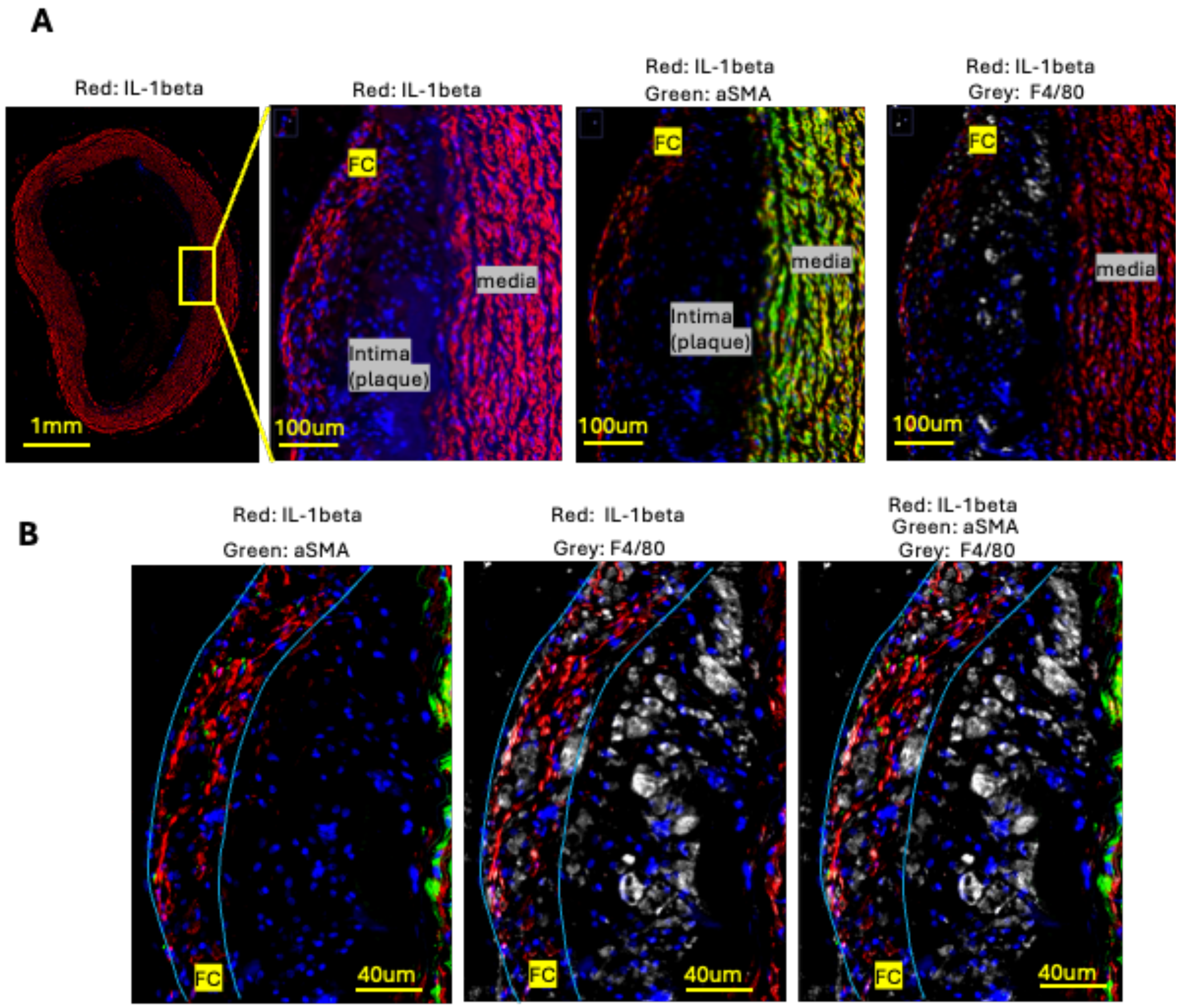


**Supplemental Figure 6. IL-1 beta immunopositivity was co-localized with vascular SMC marker in carotid media and with cells in the fibrous cap (FC) in the intima (plaque). A.** Il-1beta, aSMA (SMC marker) and F4/80 antigens were co-stained by immunohistochemistry (IHC). Low magnification images show strong IL-1 beta signal in the vascular media and in the fibrous cap. **B,** high magnification images. Il-1 beta signal in FC was co-localized with F4/80-positive macrophages. Blue curves show FC outline. Yellow bar, scale bar.

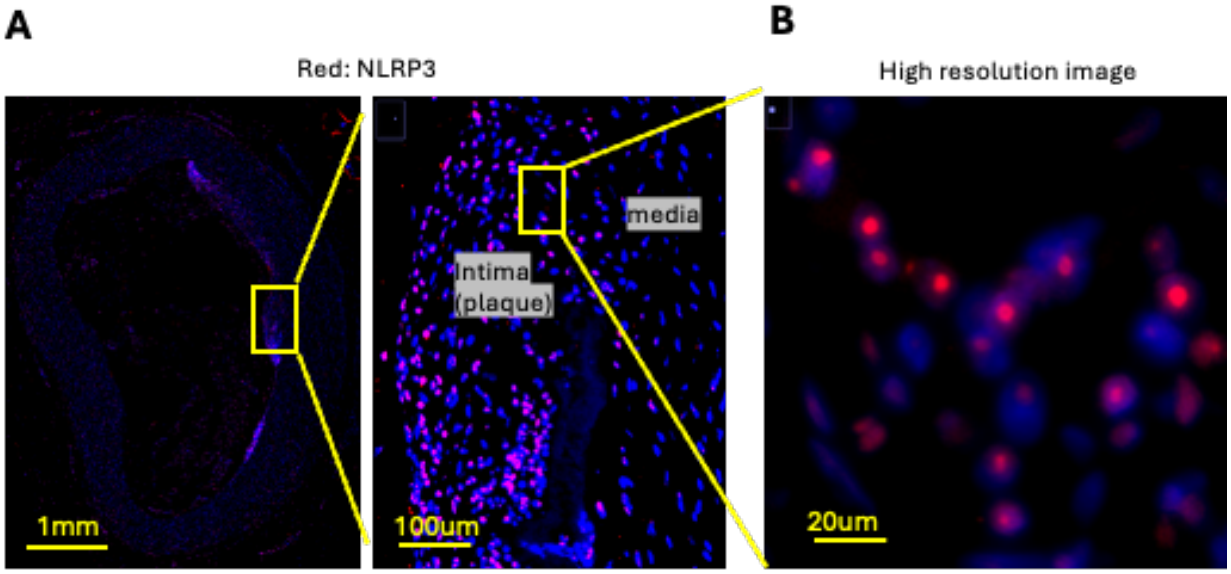


**Supplemental Figure 7. NLRP3-immunopositivity in the carotid artery section. A,** Carotid artery section was immunostained using NLRP3 antibody and co-stained with DAPI. Low resolution image shows strong NLRP3+ signal detected exclusively in plaque's cells. NLRP3+ signal in media was negligible. **B,** High resolution image show that NLRP3+ signal was mainly nuclear.
